# A Cryo-to-Liquid Phase Correlative Light Electron Microscopy Workflow for the Visualization of Biological Processes in Graphene Liquid Cells

**DOI:** 10.1101/2023.05.08.539575

**Authors:** Luco Rutten, Ben Joosten, Judith Schaart, Marit de Beer, Rona Roverts, Steffen Gräber, Willi Jahnen-Dechent, Anat Akiva, Elena Macías-Sánchez, Nico Sommerdijk

**Affiliations:** Department of Medical BioSciences, Research Institute for Medical Innovations, Radboud University Medical Center, Geert Grooteplein Zuid 28, 6525 GA Nijmegen, Netherlands; Electron Microscopy Center, Radboudumc Technology Center Microscopy, Radboud University Medical Center, Geert Grooteplein Noord 29, 6525 EZ Nijmegen, Netherlands; Helmholtz Institute for Biomedical Engineering, Biointerface Laboratory RWTH Aachen University Hospital, Pauwelsstrasse 30, 52074 Aachen, Germany; Department of Stratigraphy and Palaeontolgy, University of Granada, Avenida Fuente Nueva S/N CP: 18071, Granada, Spain

**Keywords:** liquid phase transmission electron microscopy, cryo-correlative light electron microscopy, graphene liquid cells, calciprotein particles

## Abstract

Liquid phase electron microscopy (LP-EM) has emerged as a powerful technique for *in-situ* observation of material formation in liquid. However, monitoring these processes requires the repeated interaction of the electron beam with the aqueous environment leading to the decomposition of water molecules (radiolysis), which affects the formation processes under investigation. Graphene’s ultra-conductive properties have made it an efficient way to mitigate this problem, as it acts as an electron scavenger when used as a window material. Using the strategy, the process of interest is initiated when the graphene liquid cells (GLCs) are sealed. This means that the process cannot be imaged at early time points since microscope preparation and initiation of image acquisition at the region of interest are time-consuming.

Here we report a novel cryogenic/liquid phase correlative light/electron microscopy workflow that addresses the most significant limitations of the graphene liquid cells, while combining the advantages of fluorescence and electron microscopy. This workflow allows imaging to be initiated at a predetermined space and time by vitrifying and thawing at a selected time point. We demonstrate the workflow first by observing multiple day crystallization processes and highlight its potential by observing a biological process: the complexation of calciprotein particles. With this observation, we show the exciting possibilities for LP-EM in biology.

## Introduction

Most synthetic and nearly all biological material-forming processes occur in liquid media, which severely limits the range of tools we can use to gain detailed insight into the underlying mechanisms. CryoTEM (termed cryo-EM in structural biology) provides molecular and nanoscale imaging of fast-frozen, vitrified solutions for both synthetic^1^ and biological^2^ systems. By time-resolved sample preparation it allows to study the progression of processes while keeping the sample in the native hydrated state.^3^ Nevertheless, it only provides snapshots of the specific object under observation with no information on its past or future states.

Liquid phase electron microscopy (LP-EM) has evolved as a powerful technique for the dynamic nanoscale imaging of material formation processes in solution,^4-7^ but is so far mainly applied to hard, metallic or inorganic materials.^5^ More recently, LP-EM has also been applied to dynamically imaging of soft matter systems^6,8^ and static imaging of biological^9^ and biology-derived^10^ systems in aqueous solutions. However, the dynamic monitoring of biological processes with nanoscopic details is still out of reach.^11^

The direct reason for the difficulty in imaging a dynamic biological process lies in the combination of the electron beam sensitivity of biological materials, their low electron contrast, and the necessity to repeatedly expose the system to the electron beam to monitor evolution in the process of interest. The interaction of electrons with a layer of liquid generates a complex sequence of chemical reactions (radiolysis) resulting in the formation of intermediate reactive species that influence the experimental process.^12^ However, new developments in EM technology, including the evolution of superfast, supersensitive detectors,^13^ phase plates for contrast enhancement,^14^ and sparse imaging strategies^15^ are now promising to close the gap between imaging beam sensitive materials and avoiding beam-induced damage.^8,10,16,17^ Another promising development is the use of graphene liquid cells (GLCs) for the encapsulation of samples. Here the graphene provides an ultrathin, electron-transparent viewing window^18^ and reduces electron beam-induced damage by mitigating the effect of beam-generated reactive species.^19^

For many years the use of GLCs was hampered by cumbersome preparation procedures,^20^ of which both the efficiency and reliability were low.^21^ A significant improvement in reliability was achieved through the introduction of GLC assembly by loop assisted transfer (LAT) of the upper graphene layer onto a graphene coated TEM grid.^20,21^ Another significant limitation of the use of GLCs is the offset in time (“*delay time*”) between the start of the process (typically the moment of GLC assembly), and the moment of recording the first image. This “*delay time*” includes (1) transferring the sample to the TEM, (2) finding an area of interest and (3) preparing the microscope for image acquisition.

Here, to address the *“delay time”* limitation, we take inspiration from the life sciences, where cryogenic correlative light and electron microscopy (cryoCLEM) is used to locate areas of interest with fluorescence microscopy before detailed electron microscopic investigation.^22-24^ We present a cryoCLEM investigation of GLCs that are vitrified directly after their preparation and subsequently are thawed inside the TEM for reinitiating the cryo-arrested process. Allowing (1) to minimize the *delay time* and (2) to localize areas of interest and to define the starting point (t_start_) without exposing the sample to the destructive electron beam. This cryo-to-liquid (CL) CLEM workflow is demonstrated for the investigation of the nucleation and growth of crystals. Subsequently, the CL-CLEM workflow enabled for the first time to capture a biological process. We visualize, in real time, the process that prevents ectopic calcification of our veins and tissues: the formation of calciprotein particles in cell culture medium – through the complexation of the protein Fetuin-A with excess calcium phosphate.^25^

## 2 Results

### 2.1 Workflow design

Graphene liquid cells were prepared using automated loop-assisted transfer (aLAT) on TEM grids with a 4 nm carbon support layer (Figure S1-3). The TEM grids used in this workflow are finder grids that contain compartments marked with letters and numbers that allow for the localization of areas in different imaging modalities. To easily locate the GLC liquid pockets, we add a non-reactive fluorescence dye to the sample solution during GLC sample preparation. The fluorescent signal intensity increases at the liquid pockets, where the solution is collected and confined, enabling the localization of the different pockets with cryo-fluorescence microscopy (FM) (**Figure 1**).

**Figure 1.**
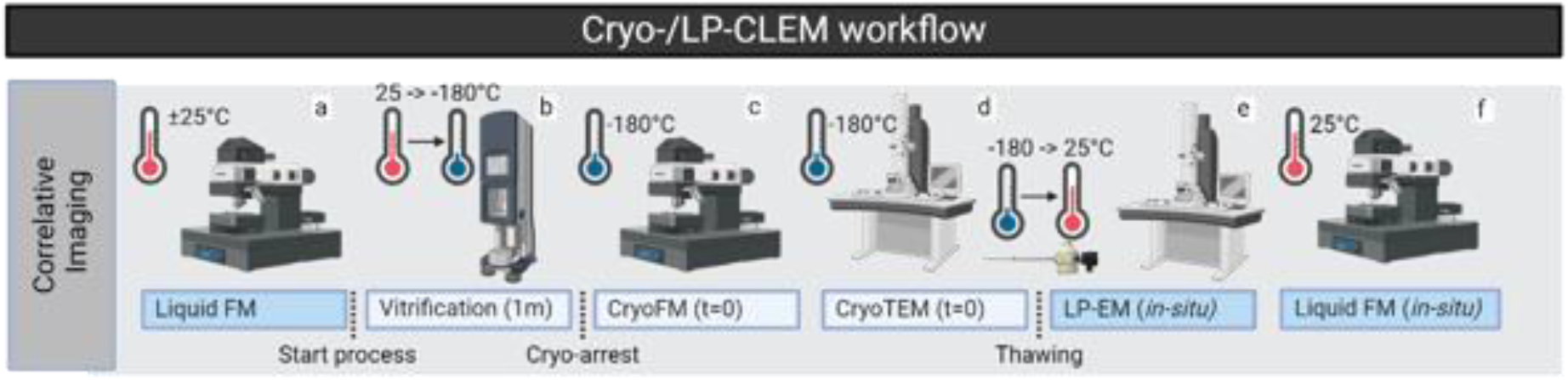
Schematic overview of the cryo-/liquid-CLEM workflow. The reaction starts once the liquid pockets are sealed, after which the reaction is cryo-arrested by plunge freeze vitrification (b). Alternatively, live fluorescence microscopy is used before vitrification to determine the optimal time of cryo-arrest (a). After cryo-arrest ROIs are located using cryo-fluorescence microscopy (c) and then imaged at nanoscale resolution with cryo-TEM (d). The process is reinitiated inside the microscope by heating-up the grid to the desired process temperature using the temperature controller of the cryo-holder, and the reaction dynamics are directly recorded using LP-EM (e). Fluorescence microscopy is used to confirm that GLCs remain intact and retain liquid after TEM observation (f).

Depending on the nature and the timescale of the process, the moment of vitrification may be chosen to be immediately at the start of the reaction or at a later time point when a more specific reaction of interest occurs. Liquid FM (called live imaging in biological studies) can be used to determine the optimal moment of vitrification and thereby to set the starting point (t_start_) for the period of interest (Figure 1a). Vitrification of the GLC-carrying TEM grid is performed through a standard protocol by plunge freezing in liquid ethane using a vitrification robot (Figure 1b). When the vitrification robot is placed in the immediate vicinity of the aLAT instrument, vitrification can be achieved within 1-2 minutes after GLC formation (t_freeze_).

After vitrification (Figure 1b), cryoFM allows localization of the GLCs without exposing the sample to electrons and to confirm the presence of frozen liquid pockets after vitrification (Figure 1c). Subsequently, the grid is transferred to a cryo-holder for TEM, after which low and higher magnification cryoTEM images are recorded of the area of interest that was selected in cryoFM (Figure 1d). In the next step, the sample is thawed by increasing the temperature to a set value (e.g. room temperature or 37° C) using the temperature controller of the cryo-holder, thereby reinitiating the process that was stopped by cryo-arrest and allowing the dynamics of the reinitialized process to be captured with LP-EM (Figure 1e). After the experiment, the grid is (again) imaged with live FM to certify the integrity of the liquid pockets during the freezing/thawing process (Figure 1f).

### 2.2 Establishing a correlative liquid-to-cryo-to-liquid workflow for targeted LP-EM

To demonstrate the potential of the proposed workflow, first a crystallization assay was used to show the ability to monitor processes at targeted positions in GLCs for multiple days. A bio-inspired crystallization solution (137 mM NaCl, 1.7 mM CaCl_2_, 9.5 mM K_2_HPO_4_, 150 µg/mL poly aspartic acid),^26^ together with a non-reactive fluorescent dye (CF®680, free acid) was encapsulated in graphene on a finder grid covered with a 4 nm amorphous carbon support film using aLAT (Figure S1). Here, the time between the application of the liquids and the closure of the GLC allowed for partial evaporation of the solution, increasing supersaturation and, thereby, the driving force for crystal formation. After encapsulation, the finder grid was immediately vitrified (t_freeze_ = 2 min) to arrest the crystallization process and imaged with cryo-confocal reflection microscopy to obtain an overview of the finder grid pattern (**Figure 2a**). CryoFM with Airyscan indicated the location of the liquid pockets on the vitrified grid (magenta spots, inset Figure 2a). The overlay of low magnification cryoTEM images with cryoFM images (100x objective, pixel size 169 nm) was used to identify and localize the hydrated liquid pockets (Figure 2b, magenta areas). In these hydrated pockets, cryoTEM imaging (pixel size 1.2 nm) showed mineral crystals that already had developed inside the GLC prior to vitrification, most probably as a result of supersaturation induced by solvent evaporation during the encapsulation process (Figure 2c).

**Figure 2.**
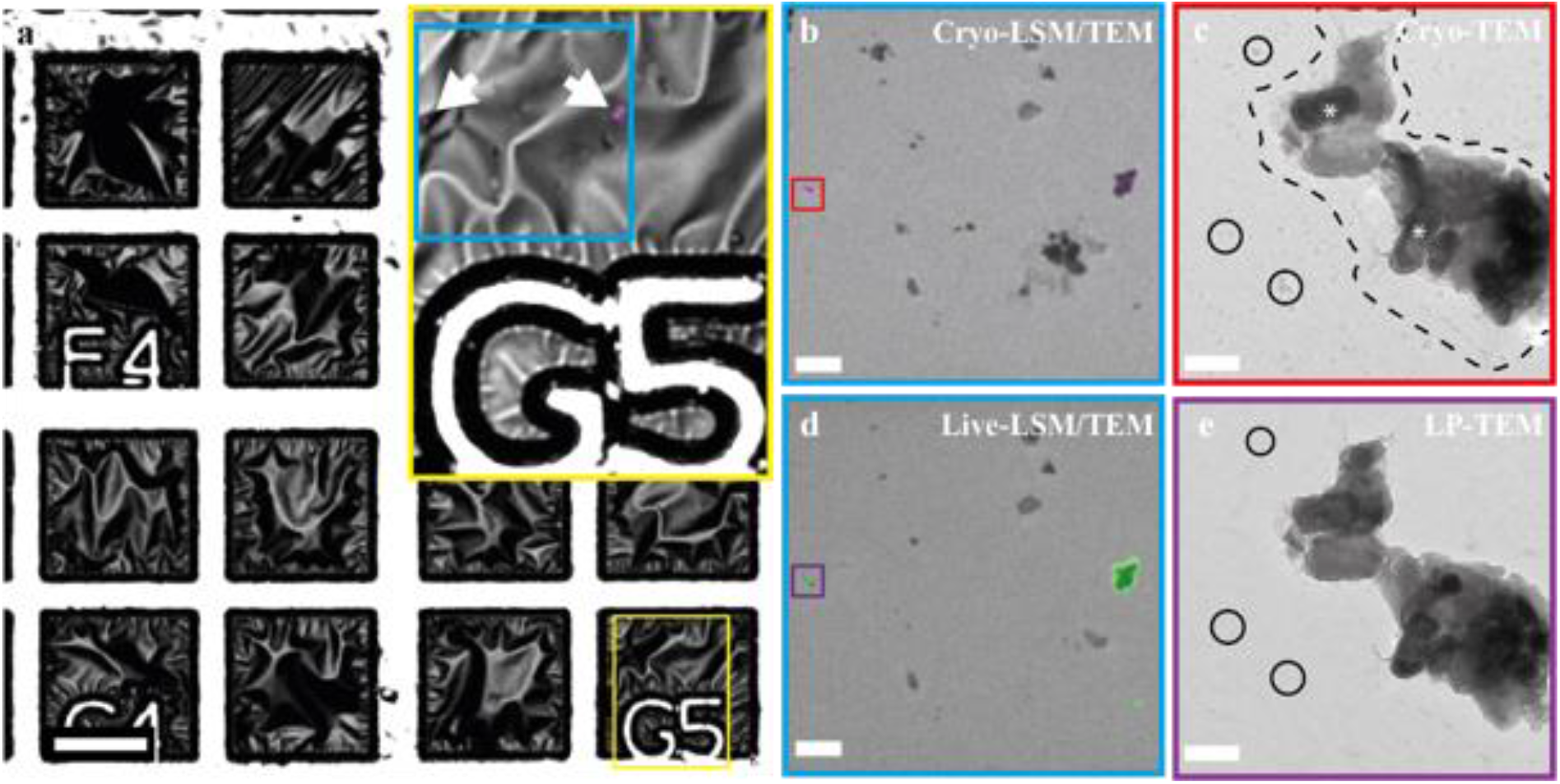
Correlative cryo-/liquid phase imaging of graphene liquid cells. a) cryoFM of the vitrified finder grid, show the overview and targeted region (G5) in cryo-confocal reflection mode. Inset shows the superimposition of the cryo-fluorescence image (magenta, arrowheads) and the confocal reflection image of the mixed solution of the buffer encapsulated in graphene and the non-reactive dye (magenta). b) an overlay of cryoFM and cryoEM image shows the enlarged image of the area marked in the blue inset (a). Magenta areas indicate the hydrated pockets. c) cryoTEM image shows the enlarged image of the area marked in (b) (red box). The crystals formed inside the liquid pockets are visible (*). The outline of the liquid pocket is indicated with a dashed line. Circles indicate ice contamination (total dose 2 e^-^/Å^2^). d) Overlay of LP-EM with liquid-FM after EM imaging (green) of the same area as in (b). Green areas indicate liquid pockets. e) Enlarged image of the area marked in (d) (purple box) showing the same liquid pocket as in (c) after thawing (total dose 4 e^-^/Å^2^). Ice crystals that were present in (c) have evaporated (empty black circles). Scale bars: a) 50 µm, b,d) 5 µm, c,e) 200 nm.

The reaction was restarted by heating the grid from –180 ° C to 20° C inside the microscope, at a rate of 5 degrees per minute, bridging between the melting point of ice and room temperature (from 0° to 20° C) in approximately 4 minutes (t_thaw_). At this time point (t_start_ = t_freeze_ + t_thaw_ = 2 + 4 min), the sample is ready for LP-EM imaging (see next section). Reaching t_start_ we were able to compare the state of the sample before and after thawing (at −170 and 20 °C, respectively, Figure 2). The images show the transformation from vitreous ice to liquid water (Figure S4), and the sublimation of the surface ice crystals (Figure 2c,e, circles) that become deposited on the sample during cryo-transfer between the FM and TEM. It is important to note that for larger pockets (> 2 µm), thawing often results in the rupture of the graphene membrane (Figure S5) and, subsequently, in the evaporation of the solution under the high vacuum of the TEM. In contrast, liquid cells with diameters between 200 and 1000nm showed excellent stability during the freezing/melting process. These liquid pockets showed fluorescence in liquid-FM (after EM imaging, Figure 2d, green areas) and remained stable for periods of up to one month (Figure S6), thus allowing the monitoring of the crystallization process over extended periods of time.

### 2.3 Monitoring Multiday Crystallization in GLCs

The presence of liquid multiple days after thawing was confirmed by superimposing low-magnification LP-TEM overviews with room temperature fluorescence images (**Figure 3a**, green areas). LP-TEM further showed the formation of rectangular and cubic crystals identified as NaCl using electron diffraction (ED, Figure S7a,b, Figure S8) and energy dispersive X-ray spectroscopy (EDX, Figure S9). Submicron-sized pockets were selected for higher-resolution monitoring of crystal growth processes (Figure 3b, Figure S7). Figure 3 also illustrates the ability to observe crystal growth processes over extended periods of time. After two days, a mineral particle is observed that, based on its shape (no facets) and the absence of diffraction contrast, represented an apparently poorly ordered (or amorphous) stage in the crystallization process (Figure 3c). After five days (Figure 3d), small domains showing diffraction contrast had nucleated from the less ordered phase. Subsequently, the two crystals coalesced, likely by Ostwald ripening, and transformed into a cubic NaCl crystal after seven days (Figure 3e).

**Figure 3.**
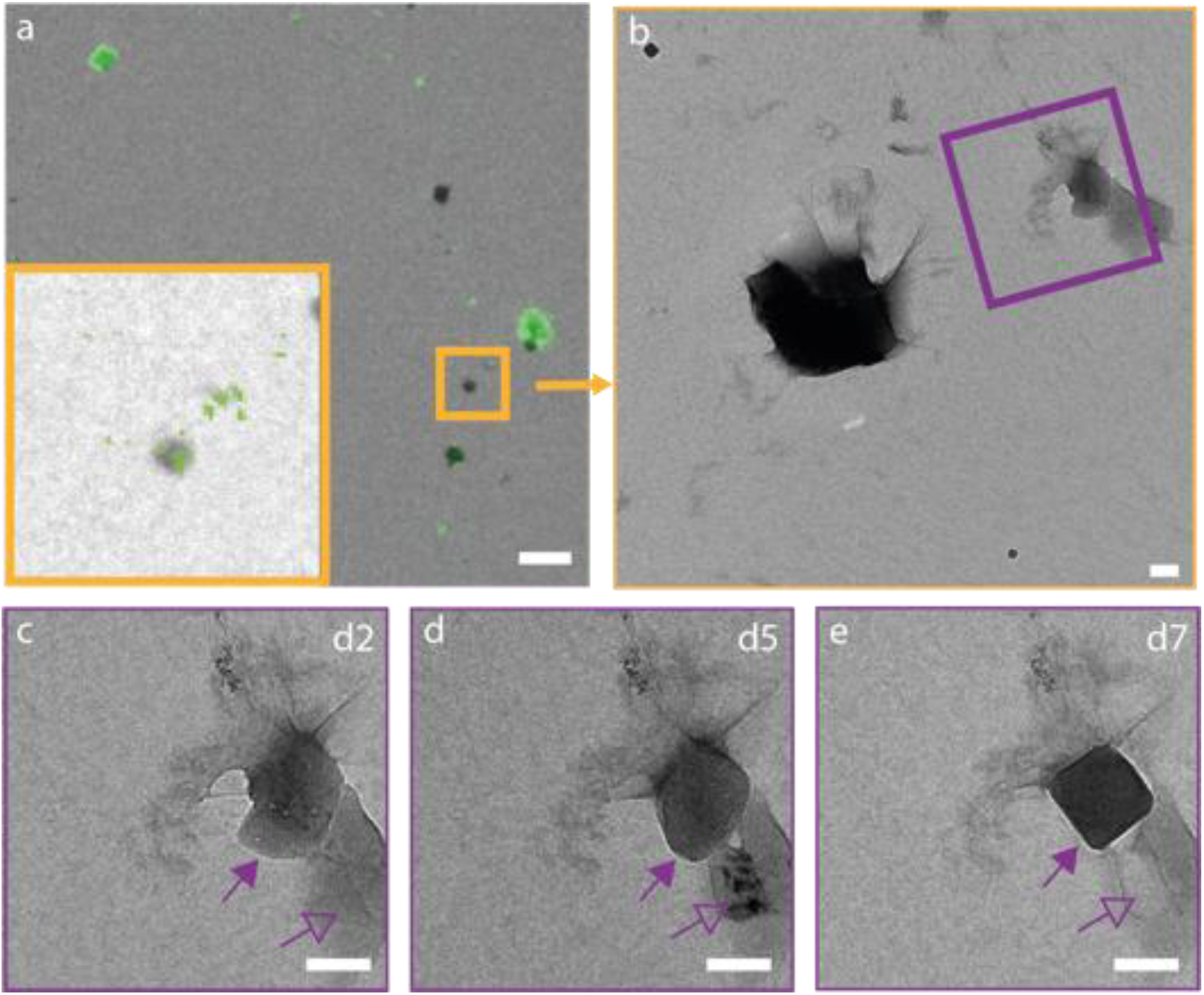
LP-EM visualization of crystallization processes inside a GLC. a) LP-EM overview recorded two days after thawing, overlaid with liquid-FM (green) to indicate the GLCs. Inset shows a high-magnification LP-EM image of the GLC in the orange box. b) LP-EM of the area marked by the orange box in (a) where multiple GLCs are present. c) Two different but interconnected GLCs (purple box in (b)) imaged two days after thawing (open and closed purple arrows). d) Condensation (solid arrow) and phase transformation (open purple arrow) observed within the pockets five days after thawing. e) Cubic NaCl crystal resulting from an Ostwald ripening process observed seven days after thawing. Scale bars: a) 5 µm, b) 500 nm, c-e) 200 nm. Accumulative dose: a) 0.75e^-^/Å^2^; c) 0.15 e^-^/Å^2^; d) 0.30 e^-^/Å^2^; e) 0.45 e^-^/Å^2^

The liquid nature of the crystal’s environment was confirmed by the visible outline of the liquid pockets, the high contrast graphene wrinkles around the crystal, and the intensity gradient surrounding the crystal (Figure S7d).^27^ Further proof for the presence of liquid came from exposure of the pockets to a high electron dose (total electron dose > 100 e^-^/Å^2^), which resulted in direct bubbling of the liquid due to radiolysis (Figure S10). Crystals that developed on parts of the grid that were not encapsulated by graphene showed no crystallization processes (Figure S11).

### 2.4 Monitoring biological self-assembly in GLCs

After developing this GLC-based CL-CLEM workflow for inorganic crystallization processes, we set out to demonstrate its application for biological systems. For this we selected the complexation of calcium phosphate with the protein fetuin-A, an important process in the blood stream to prevent ectopic mineralization.^28^ When calcium phosphate levels in the blood exceed physiological concentrations, monomeric fetuin-calcium phosphate complexes (calciprotein monomers, CPMs), which are cleared by the kidneys.^29^ In pathological situations (e.g. chronic kidney disease) this clearance mechanism fails, causing CPMs to coalesce into amorphous spherical aggregates called primary calciprotein particles (CPPs), and ultimately into crystalline secondary CPP that are cleared by endothelial cells and phagocytes, respectively^30^ and have been suggested to contribute to (cardio)vascular calcification.^31^

To study the mechanism of CPP formation we used the complexation of calcium phosphate in standard cell culture medium (minimum essential medium, α-MEM) containing 10% fetal calf serum (FCS) by the native fetuin-A present in the FCS. 3.50 mM phosphate was added to the medium to create a biomimetic supersaturation,^32^ as well as 1 mol% CF®568-labeled fetuin-A to identify the formed CPPs with FM. CF®680 was added for the fluorescence identification of liquid pockets. The mixture containing 3.66 mM phosphate, 1.73 mM calcium, 2.64 mg/mL fetuin-A (determined by Western Blot, not shown) was placed on the grid and enclosed in GLCs by aLAT. Dynamic light scattering (DLS) showed in bulk reaction that with these concentrations of reagents CPP formation occurs within 36 minutes (Figure S12).

As biological self-assembly processes are sensitive to concentration and temperature, the aLAT instrument was adapted to work at controlled humidity and temperature (Figure S13). This allowed us to study the formation of CPPs under near-native conditions, preparing samples at 37 °C and 96% relative humidity, i.e., at body temperature and at the native fetuin-A concentration in fetal blood.^33^ Directly after encapsulation, the grid was vitrified (t_freeze_ = 2 min) and subsequently imaged using cryoFM and cryoTEM to identify the regions of interest (Figure S14).

The vitrified grid was first slowly (0.5 degrees/min) heated to 0 °C and then quickly (20 degrees/min) heated up to 37 °C (t_thaw_ = 2 min). At this point (t_start_ = 4 min), image acquisition was started by applying a very low electron dose (applied dose: 0.05e^-^/Å^2^/ image; image interval 60-120 sec). A region of interest was chosen within a GLC (**Figure 4a**) to monitor CPP formation over a period of 11 minutes (Figure 4d-h, Figure S15), overlapping with the period investigated with DLS (Figure S12).

**Figure 4.**
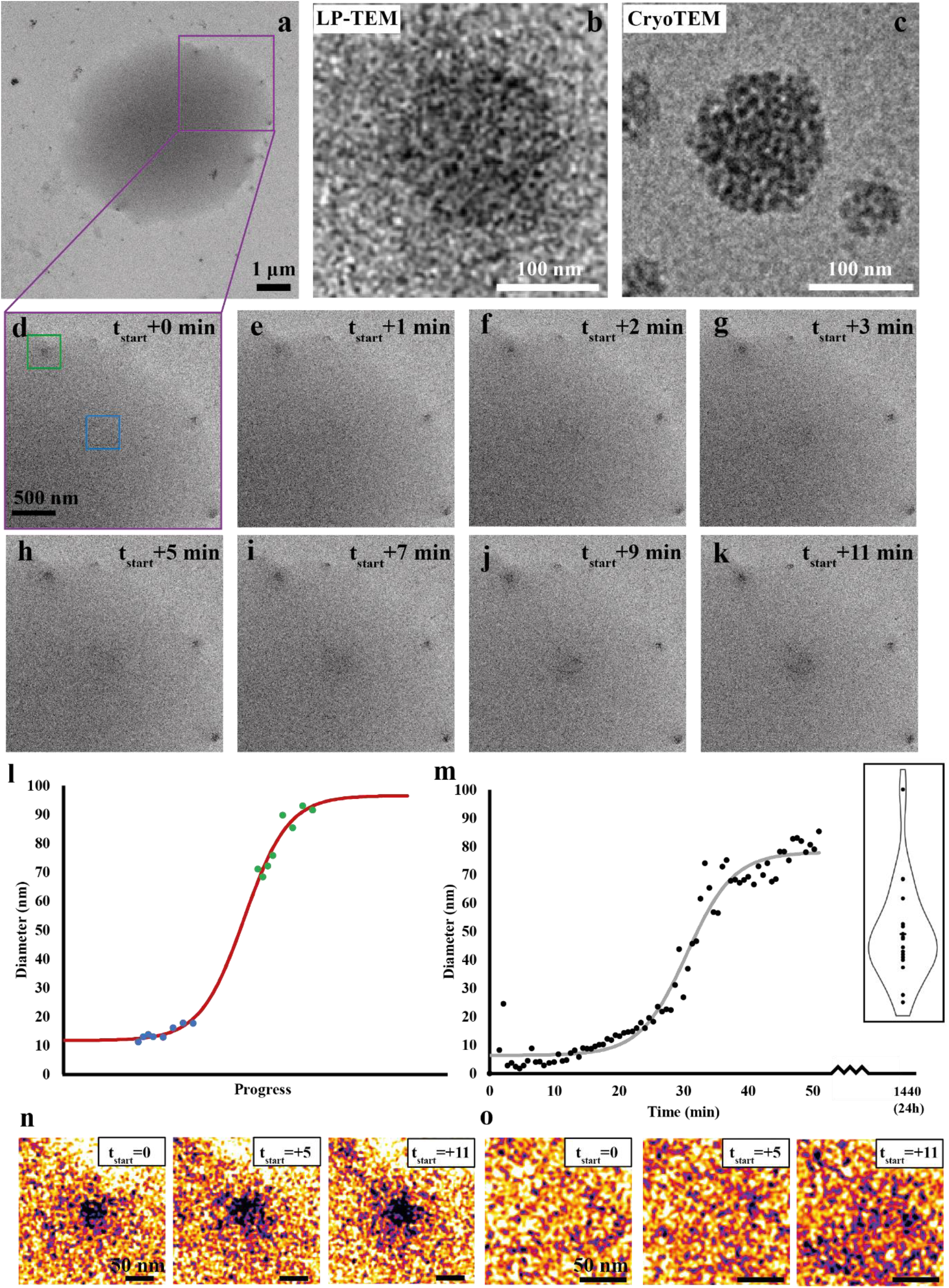
Formation of calciprotein particles in a graphene liquid cell using the cryo/LP-CLEM workflow. a) Overview of the graphene liquid cell. b) Mature CPP formed in a bulk solution and imaged with cryoTEM. c) CPP after 17 minutes of formation in a GLC. d-k) Zoom-in of magenta square area in (a) were two CPP formation processes are visible over a period of 11 minutes. Green square indicates the region where a 40 nm CPP is already visible. In the blue square only CPMs are visible. l) Analysis of the particle size in the blue and green region. Red line fits a sigmoidal growth. m) CPP formation monitored with dynamic light scattering showing a typical S-curve. Plot shows maximum intensity derived from data in Figure S15 n) CPP size distribution after 24 hours determined with cryoTEM. o) Analysis of the diameter growth of the CPP in the green square. Corresponding images in white/red contrast are visible at timepoints 1, 5 and 11 minutes. p) Analysis of the diameter growth of the CPP in the blue square. Corresponding images in white/red contrast are visible at timepoints 1, 5 and 11 minutes. Scale bar: a: 1 µm, b,c: 100 nm d-k: 500 nm, o,p: 50 nm. Dose d-k: 0.05e-/Å^2^/ image.

Calciprotein monomers (CPMs) and small CPPs were both already present at the first time point (t_start_ = 4 min) (Figure 4a, Figure 4d, Figure 4n,o). Within the region of interest, we observed both the nucleation of new and the growth of already existing CPPs (Figure 4d-4k). Image processing using denoising, contrast enhancement and shading correction, revealed that nucleation (Figure 4a, blue square; Figure 4o) involved the gradual densification of the liquid by the gathering and clustering of CPMs and their aggregation into early-stage CPPs (t=t_start_ +11 min, diameter ∼18 nm). In a neighboring region, the development was observed (Figure 4a green square, Figure 4n) from a smaller (∼70nm) to a more mature CPP (∼90 nm). During this process, a ring containing single CPMs surrounded the CPP (Figure 4n), suggesting growth occurred by particle absorption.

To assure that our observations were not simply the result of interactions of the solution with the electron beam, we compared both the reaction kinetics and the final result with bulk reactions. CryoTEM revealed that particles present in bulk solution after 24 hours had similar morphology (Figure 4c) and size distribution (50-100 nm) (Figure 4m, inset) that agreed well with the diameter derived from DLS of mature sample (Figure 4m) and the mature particles observed in LP-EM (∼90 nm, Figure 4b,l), which also agrees with literature data. Analysis of the DLS data showed that the maximum signal intensity built up to a plateau at 80 ± 3 nm over the course of 50 min, in line with our LP-EM observations. Specifically, the DLS data followed an “S”-curve (Figure 4m) and hence points to an autocatalytic assembly process.^34^ Although we cannot directly compare collective and single particle data, the development of particle size in our LP-EM experiments could be plotted on a similar S-curve, suggesting a similar process kinetics as observed in solution (Fig 4l).

## 3. Discussion

LP-EM uniquely provides the opportunity to observe the dynamics of materials formation at the nanometer scale and, hence, to observe the very first moments of these processes, which are often determinants of the outcome. This requires knowing where to find the relevant regions for imaging inside the TEM and being able to initiate the process of interest in a moment of choice inside the microscope.

By encapsulating the reaction medium in GLCs and immediate vitrification of the grid, our CL-CLEM workflow enables the locating of areas of interest using fluorescence microcopy and subsequently imaging the start of the reaction with LP-EM by reinitiating the cryo-arrested processes through sample thawing, irrespective of the material under investigation.

In recent years, researchers have demonstrated the live imaging of soft matter processes using LP-EM, such as the formation of polymeric vesicles^6^ or the fusion of micelles^35^ in commercially available flow-cell holders. However, to date, the electron dose and resolution requirements have prohibited the detailed investigation of biological processes.^8^ Nevertheless, the observation of processes involving biological materials has recently been reported using GLCs, but so far, only by triggering the process with the electron beam^36^ or after a *delay time* (*delay time* > 20 min) required for the assembly and transfer of the sample.^37^

In traditional liquid cell holders, a process or reaction can be conveniently started by flowing in solutions or liquids that initiate the process of interest during the observations, catching t=0 in materials formation processes also strongly depends on imaging at the right location at the right time. In GLCs, on the other hand, the solution composition is essentially defined upon the closure of the liquid pockets, which significantly limits the way processes can be triggered after the GLC is prepared. Currently this has only been demonstrated through triggering processes with the electron beam, where this method is restricted to the initiation of metal or inorganic nanoparticle formation,^38^ or their dissolution by etching.^39^

GLCs in principle are suitable for studying processes involving temperature-responsive components, such as the temperature-triggered morphology changes of polymer nanoparticles that so far was only demonstrated using classical flow cells.^40^ However, for processes not being triggered inside the microscope, a “*delay time*”, the time from encapsulation to recording the first LP-EM image – is unavoidable. The *delay time* for finding a pocket without using our workflow is in our hands in the order of 20-30 minutes, but without the use of FM this procedure does not guarantee the presence of liquid in that pocket. Moving to a next potentially interesting pocket will further increase the *delay time*.

The ability to locate GLCs with FM, i.e., without exposure to the electron beam prior to capturing the process-of-interest, is an important asset in imaging beam-sensitive materials. Where fluorescence is nowadays routinely used to locate regions of interest in cryoCLEM^8^ for cryo-electron tomography, its use in navigating samples for LP-EM is virtually unexplored. So far, the use of FM in combination with LP-EM has been used to locate water-containing GLCs,^21^ to confirm the hydrated state of fixated cells after TEM imaging^9^ and to investigate electron beam-induced reactions.^41^ Our CL-CLEM approach not only offers the possibility to locate the GLCs with FM in a cryogenic state prior to the start of the process but also to confirm the regaining of the liquid state in the dynamic imaging part of the workflow. With regard to the integrity of biological samples during the above experimental sequence, it is important to note that proteins can reliably maintain their molecular structure during laser-induced freezing-thawing-refreezing cycles.^42,43^

The CL-CLEM workflow presented above allows targeting of specific pre-selected areas to re-start previously cryo-arrested processes and to image them from the moment of re-initiation so that the very early stages can be studied with great detail. Our work shows that in this way, we can capture biological processes in GLCs by imaging at a low dose (0.05 e^-^ /Å^2^/image, total dose 0.4 e^-^/Å^2^) and at nanometer resolution (2.67 nm/pixel), providing us the necessary details to follow a process. Here, the biological process we focused on was CPP formation: CPP crystallization and precipitation can lead to vascular calcification, which is a major health issue. The ability to visualize biological processes in high resolution will aid in understanding the mechanism and can be used to develop therapeutic strategies. Besides, it highlights the possibility to investigate the morphological dynamics of other biological materials such as vesicles, proteins, and membranes.

Imaging at relatively low electron doses (dose rate 2-5 e^-^ Å ^-2^ s^-1^, total dose 100-900 e^-^ Å ^-2^) of a biological process in GLCs was demonstrated with high resolution (at 0.45-0.74 nm/pixel).^37^ In the present work the accumulative dose used (0.5 e^-^ Å ^-2^) to study the CPP formation is a magnitude lower than the safe dose (5 e^-^ Å ^-2^) we established prior to the experiment. This means it would still be possible to either increase temporal resolution (by increasing the number of recorded images), and/or increase the resolution (by decreasing the pixel size). However, it is important to place these results in the context of the general dose requirements for imaging biological processes. While dynamic imaging of morphological changes in biological-derived systems is possible using an electron dose of 30 e^-^ Å^-2^,^36^ enzymes already start losing their activity at much lower electron doses, with reported critical accumulative dose around 10^−4^ e^-^ Å^-2.44^

Although this dose limitation in LP-EM will still allow to imaging morphological dynamics that do not directly rely on enzymatic activity, it underlines the need to rationally design imaging strategies for the live/in situ observation of complex biological processes. Moreover, it implies that we should refrain from being guided by electron doses typically used in cryo(T)EM as prevention of loss of observable damage does not exclude alteration of biological activity.

## 4. Conclusion

This paper introduces a novel cryo-to-liquid CLEM workflow that comprises (1) plunge freezing a sample at a predetermined time point, (2) localizing the region of intertest without the exposure of electrons using FM, and (3) reinitializing the reaction by thawing the vitreous sample in the microscope during *in-situ* imaging. This CL-CLEM workflow allows for the observation of beam-sensitive biological processes by using graphene as window material. A recent review already envisioned that the use of LP-EM to capture biological processes could solve grand challenges in biology, leading to new clues in biomedical sciences and improving therapeutic strategies.^8^ However, for this, controlling radiation damage, conducting control experiments, and choosing the optimal experimental settings were considered key factors. Our observation of the complexation of fetuin-A with calcium/phosphate, under biological relevant conditions, shows that LP-EM indeed can be a powerful tool in biology and biomedical research following careful experimental design.

## 5. Experimental Section/Methods

### Graphene encapsulation

Gold 200 mesh ultrathin carbon film finder grids (CF200F1-AU-UL, Electron Microscopy Sciences) were encapsulated with graphene by automated loop-transfer using a Naiad-1 system (VitroTEM, Delft, The Netherlands). Copper chips coated with graphene by CVD were acquired (VitroTEM, Delft, The Netherlands) and placed inside a Naiad-1 (VitroTEM, Delft, The Netherlands). The machine etches away the copper with 0.2 M ammonium persulfate and washes the graphene with Milli-Q. CPPs were encapsulated with the modified NAIAD-1 Mod-T (Fig. S13).

### Crystallization essay

After washing the Milli-Q is replaced with the crystallization buffer (0.85x PBS, 1.7 mM CaCl_2_, 9.5 mM K_2_HPO_4_, 150 µg/mL poly aspartic acid). A finder grid is placed in the NAIAD instrument and a 3 µL droplet of crystallization buffer with 0.1 mg/mL CF®680 dye (Biotium, Fremont, CA, USA) is added on the grid. The graphene is transferred together with the crystallization buffer by automated assisted loop transfer to the grid and the excess liquid is blotted away, which results in the sealing of the liquid pockets.

### Calciprotein particle complexation

Fetuin-A/mineral complexes were prepared in a solution of 10% fetal calf serum (FCS) in alpha minimal essential medium (αMEM) (total volume 1 mL). The solution was supplemented with NaH_2_PO_4_ (2.71µL, 0.5M stock) and Na_2_HPO_4_ (4.28µL, 0.5M stock) and direct applied on the grid.

### Cryo-LSM

Encapsulated grids were vitrified by plunge freezing in liquid ethane using a Vitrobot Mark IV (Thermo Fisher Scientific, Eindhoven, The Netherlands) at room temperature. After vitrification, the TEM grids were loaded into a universal TEM cryo-holder (349559-8100-010, Quorum technologies, Laughton, UK) using the ZEISS Correlative Cryo Workflow solution, which fit into the PrepDek® (PP3010Z, Quorum technologies, Laughton, UK). The universal cryo-holder containing the samples was transferred into an adapter to fit the LM cryo-stage (CMS-196, Linkam scientific inc.), which was used with an upright confocal laser scanning microscope (LSM 900, Zeiss microscopy GmbH) equipped with an Airyscan 2 detector.

Reflection microscopy and fluorescence microscopy were recorded in three different resolution steps. First, overview images at low resolution were made (C Epiplan-Apochromat 5x/0.2 DIC, Zeiss; pixel size 520 nm) using reflection microscopy (0.02% 640nm laser with 500V master gain; Detection 549 - 700) and fluorescence microscopy (4% 640nm laser with 900V master gain; Detection 450 - 700), to visualize the TEM grid bars and liquid pockets, respectively. Next, medium resolution images (C Epiplan-Apochromat 10x/0.4 DIC; pixel size x-y: 195nm) were recorded from regions of interest (ROIs), on various areas of the grid, with the same excitation/emission settings as for the low resolution. Finally, the ROIs were recorded at high resolution (C Epiplan-Neofluar 100x/0.75 DIC; pixel size x-y: 104nm). Settings had to be adjusted for this 100x objective (Reflection microscopy: 0.2% 640nm laser with 500V master gain; Detection 549 – 700 and Fluorescence microscopy: 5% 640nm laser with 950V master gain; Detection 450 - 700). All the fluorescent images were recorded in the super-resolution mode and were processed afterwards using the auto filter in Zen Blue. The images were displayed in Zen connect, a plugin the in Zen blue software (Zeiss system software). Within this program, all the recorded images are stored with its coordinate, allowing navigation with high resolution objectives.

### Live-FM

The Zen connect file was opened which stored the previous recorded cryo LM images. The grid was placed on an objective glass and imaged with an upright confocal laser scanning microscope (LSM 900, Zeiss microscopy GmbH) equipped with an Airyscan 2 detector. The images were recorded in three different resolution steps. First, an overview image at low resolution was made C Epiplan-Apochromat 5x/0.2 DIC, Zeiss; pixel size 624 nm) using reflection microscopy (0.02% 640nm laser with 550V master gain; Detection 549 - 700) and fluorescence microscopy (2% 640nm laser with 900V master gain; Detection 450 - 700). This image was used to align previous cryo-session for easy navigation to the ROIs images with high resolution in cryo. Next, a medium resolution was recorded from the ROIs (C Epiplan-Apochromat 10x/0.4 DIC; pixel size x-y: 195nm), followed by a higher resolution image of each ROI (Plan-Apochromat 20x/0.8 M27, Zeiss; Pixel size 98nm). The settings for the medium and higher resolution were the same as for the low resolution. All the fluorescent images were recorded in the super-resolution mode and were processed afterwards using the auto filter in Zen Blue.

### Cryo-/LP-TEM imaging

Images of the CPP complexation were acquired with a TALOS (Thermo Fisher, Eindhoven) equipped with a Falcon 4i direct electron detector operated at 200 kV. Images of the crystallization were acquired with a JEOL JEM 2100 (Jeol Ltd., Tokyo, Japan) equipped with a Gatan 890 Ultrascan (Pleasanton, CA, USA) operated at 200 kV using low dose software (Serial EM). EDX mapping was performed with a JEOL JEM 1400 (Jeol Ltd., Tokyo, Japan) equipped with a JED2300T (Jeol Ltd., Tokyo, Japan) and Tungsten filament operated at 60 kV.

### Heating-up

Vitrified grids were placed inside a Gatan 914 high tilt cryo-holder (Pleasanton, CA, USA). The liquid nitrogen was removed from cryo-transfer holder without removing the holder from the microscope. The temperature controller unit was used to heat-up the tip of the holder (0.75 mA) in approximately two hours to 0 °C and then quickly (2 min) to room temperature or 37 °C.

### Image analysis

Processing of LP-EM images was done in Fiji. The shading correction was done with a bandpass filter of 100 and a Gaussian blur with sigma 1 was used for denoising.

### Dynamic light scattering

DLS measurements were performed on the Zetasizer Pro (timelapse). The fetuin-A/mineral complex solution was measured directly after production of the particles to determine particle size. For timelapse measurements, 10% FCS in αMEM was measured at 37°C before the additional phosphate was added. After phosphate addition, measurements were taken approximately every 90 seconds during the first 40 minutes, while incubating the solution in the Zetasizer at 37°C.

## Acknowledgements

LR, JS, BJ, MdB, AA, RR and NS are supported by the European Research Council (ERC) Advanced Investigator grant (H2020-ERC-2017-ADV-788982-COLMIN). EMS is supported by Marie Curie Individual Fellowship, MSCA-IF-2020 DYNAMIN (101031624), the Research Program Juan de la Cierva Incorporación (IJC2020-043639-I) funded by MCIN/AEI/10.13039/501100011033 and European Union NextGenerationEU/PRTR, and the project PID2022-141993NA-I00 funded by MICIU/AEI/10.13039/501100011033 and ERDF/UE. WJD is funded by the Deutsche Forschungsgemeinschaft (DFG, German Research Foundation) – TRR 219 – Project-ID 322900939, and 403041552.

