## Supporting Information for "A Cryo-to-Liquid Phase Correlative Light Electron Microscopy Workflow for the Visualization of Biological Processes in Graphene Liquid Cells"

**Automated loop-assisted transfer**

A reliable method for the transfer and formation of liquid cells with graphene windows is essential for LP-TEM experiments. This resulted in the fabrication of multiple GLC designs such as graphene-on-graphene pockets, graphene sealed nano-wells and liquid flowing cells[44]. However, finding a cheap, functional, accessible and reliable method to fabricate GLCs remains difficult[45]. Loop-assisted transfer (LAT) is often the method of choice to transfer graphene[21,46,47], but it still involves a highly delicate manual handling where the integrity of graphene is easily lost[20].

The aLAT GLC assembly (Figure S1) provided a significant increase in both the number (±5x) and size of the liquid pockets compared to manual assembly[21] (Figure S2a). Each 90 $\mu$m x 90 $\mu$m grid square contained on average 20 ± 8 GLCs (Figure S2a) with 60% being between 0-300.000 $nm^2$ (0.3 $\mu m^2$) in area while 10% of the liquid pockets were larger than 500,000 $nm^2$ (0.5 $\mu m^2$) (Figure S2b), equivalent to circular liquid pockets with diameters larger than 400 and 800 nm, respectively. The size of the liquid pockets was determined by automated image analysis (Figure S3).

The automated loop-assisted transfer (aLAT) has been shown to outperform the manual method in both number of cells produced and cell size. aLAT yield approximately a 5x higher amount of GLCs,[21] and the size of the liquid cells is 2 times higher when compared with manual transfer (ranging from ±100 $nm^2$ to over 6.000.000 $nm^2$). The optimal size of a GLC will ultimately depend on the process of interest. The highest resolution can be obtained in smaller liquid cells, where the liquid layer thickness is minimized. However, when a larger volume is required, bigger GLCs are necessary.

Additionally, the aLAT method offers a high degree of repeatability in terms of the volume of liquid transferred. It should be noted that the initial solution deposited on the grid will mix with the solution transferred by the loop. Since the collection of the graphene is automated, the amount of solution in the loop is consistent (±1 µL), which is an essential parameter for calculating the final concentration of the encapsulated solution that will finally trigger the reaction.

Supporting the graphene by encapsulated material or a patterned framework [48,49] could enlarge the encapsulated volume. Since GLCs are sealed volumes, the absence of flow prevents the ability to change or introduce reactants during the process. The mitigation of these limitations could involve the design of liquid-flowing graphene chips[53] that allow the vitrification and correlation of fluorescence microscopy to cryo-/LP-EM.

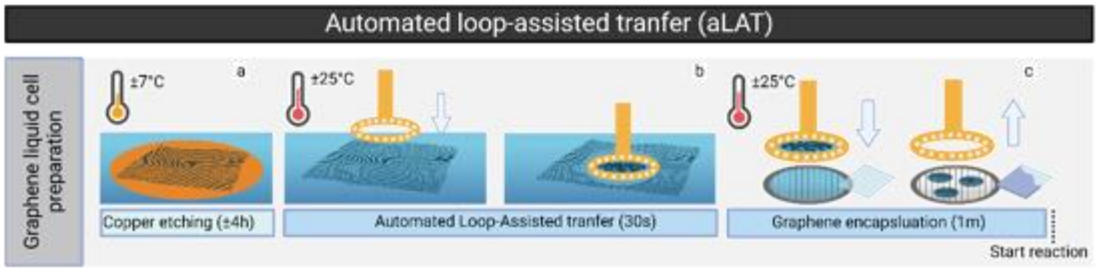


**Figure S1. Workflow for automated loop-assisted transfer (aLAT)**. The GLC preparation starts with the etching of a cupper chip supporting the graphene (Figure S1a). Thereafter the graphene is picked up by a loop and the liquid in the loop is carrying the graphene (Figure S1b) to the grid. Here the liquid in the loop and the liquid on the grid are mixed (Figure 1Aiii)[21] and subsequently excess liquid is blotted away (Figure S1c), upon which the GLCs are formed.

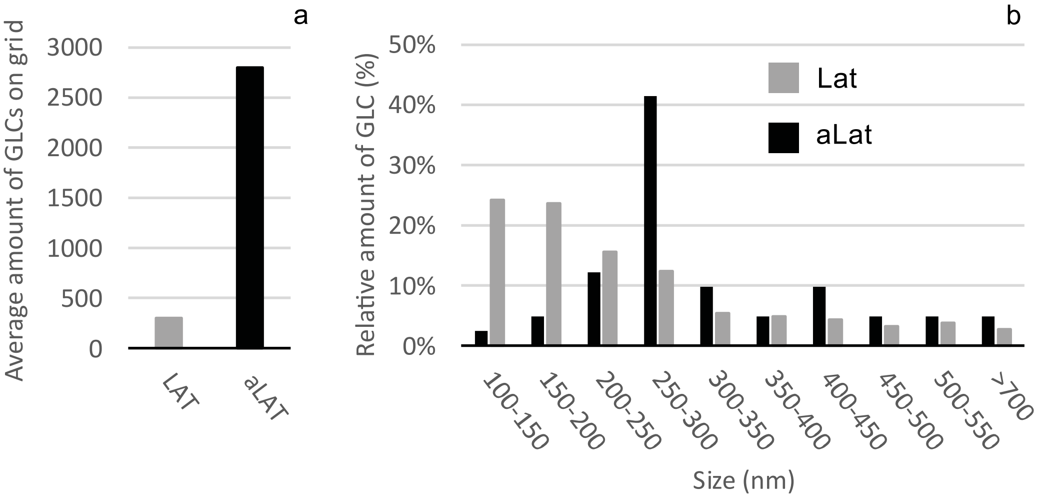


**Figure S2. Throughput comparison between loop-assisted transfer (LAT)[21] and automated loop-assisted transfer (aLAT) preparation methods.** a) Average number of

GLCs formed on a TEM grid with both methods (extrapolated from several grid squares). b) Size distribution of the viable graphene liquid pockets after grid thawing, based on the correlation between live-FM and LP-TEM and calculated by automated image analysis (Fiji).

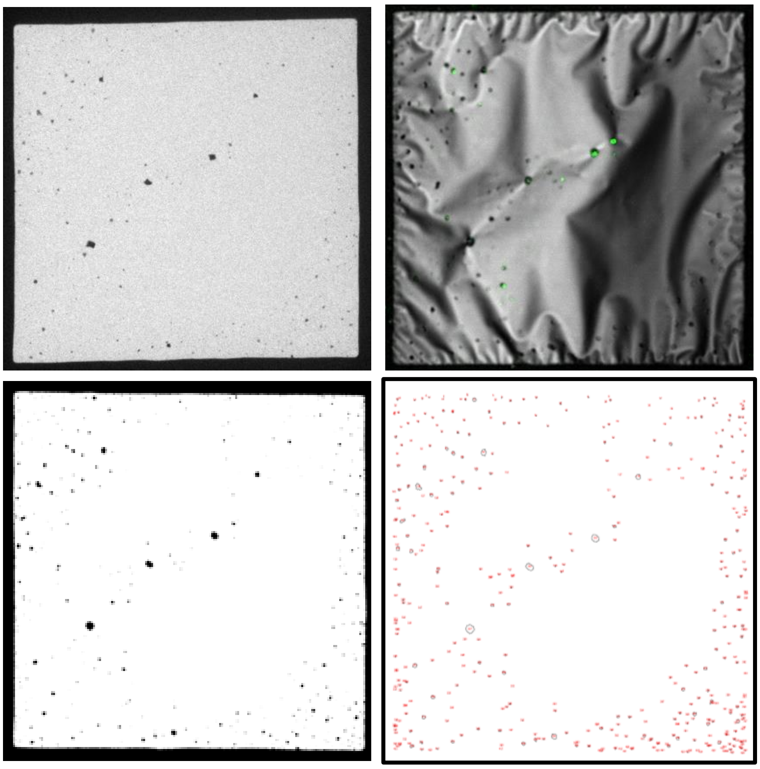

**Figure S3: Automated size/amount of liquid pockets determination.** a) Overview cryo-TEM image. b) Live-fluorescent image. c) Thresholding of the cryo-TEM overview image. d) Automated determination of size and amount of liquid pocket using Fiji. Only the pockets that showed a fluorescent signal were used for statistical determination of the size and amount of GLCs.

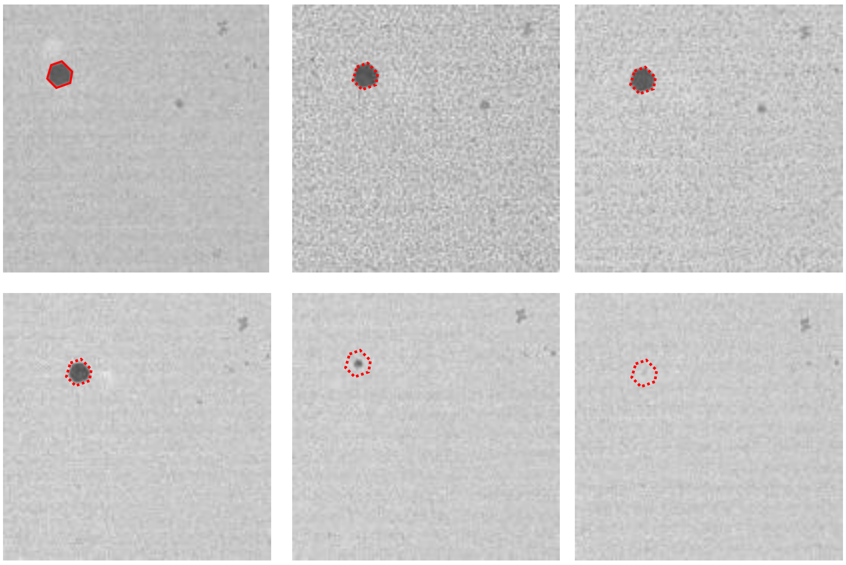

**Figure S4:** Increasing the temperature of the TEM holder inside the microscope from -180 °C to room temperature results in a phase transformation of the vitreous ice and melting of the large hexagonal ice particle (encircled dotted line).

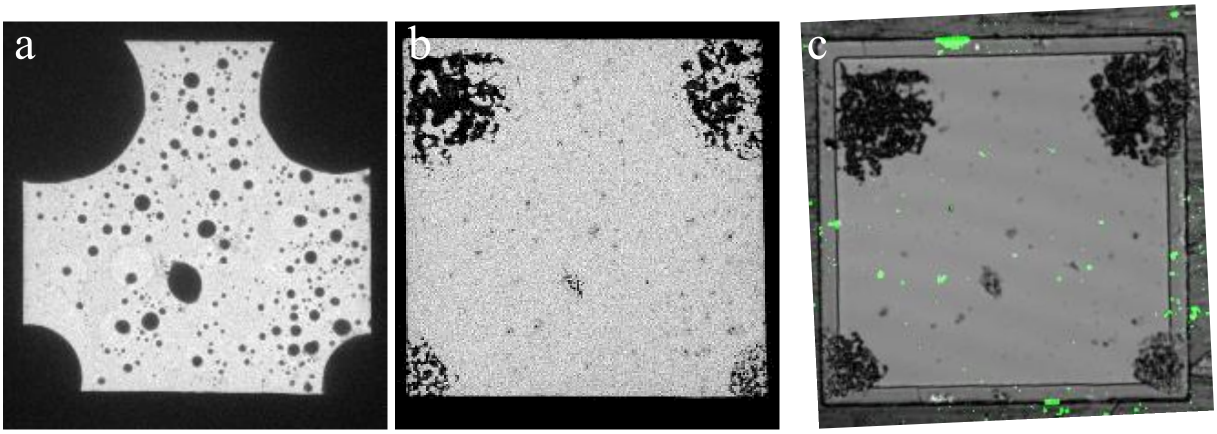


**Figure S5:** Collapsing of larger liquid pockets after thawing. a) Cryo-TEM overview showing multiple large liquid pockets. b) TEM overview of the same region directly after thawing the grid. Many large liquid pockets are collapsed. c) Live-FM of same area showing which GLC are still hydrated.

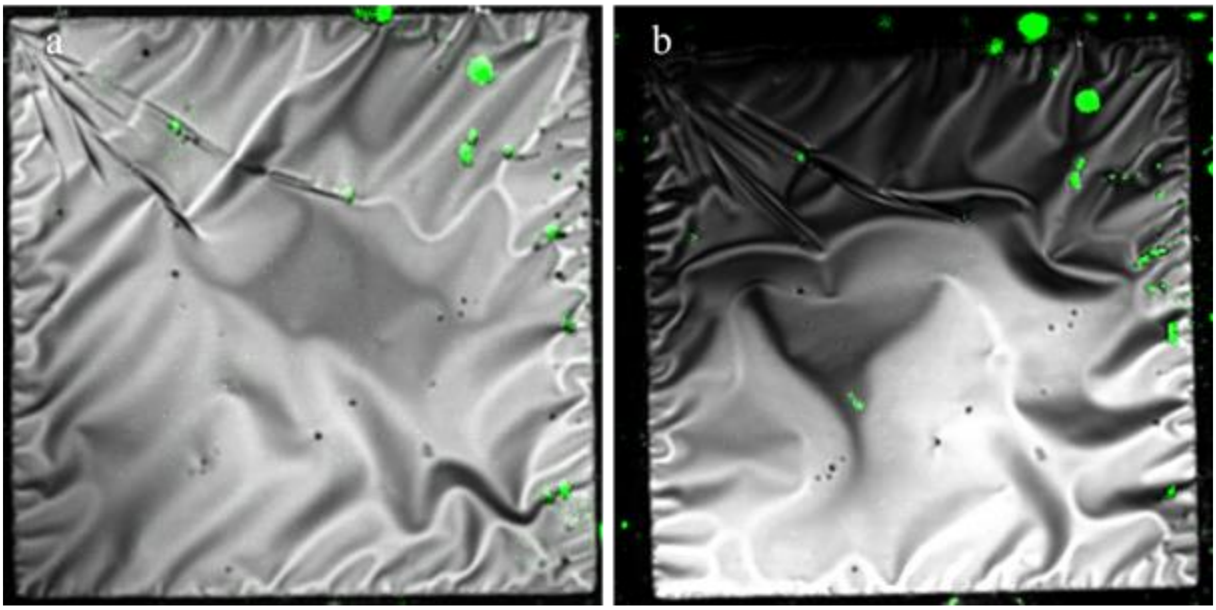


**Figure S6:** Live-FM of the tagged GLC (green) and confocal reflection microscopy (grays) 1 day (a) and 1 month after thawing (b) show that some GLC still contain liquid one month after thawing.

**Demonstrating the liquid in the pockets:**

The contrast difference surrounding the crystals marks the outline of the liquid pocket (Figure 3c, dashed line).

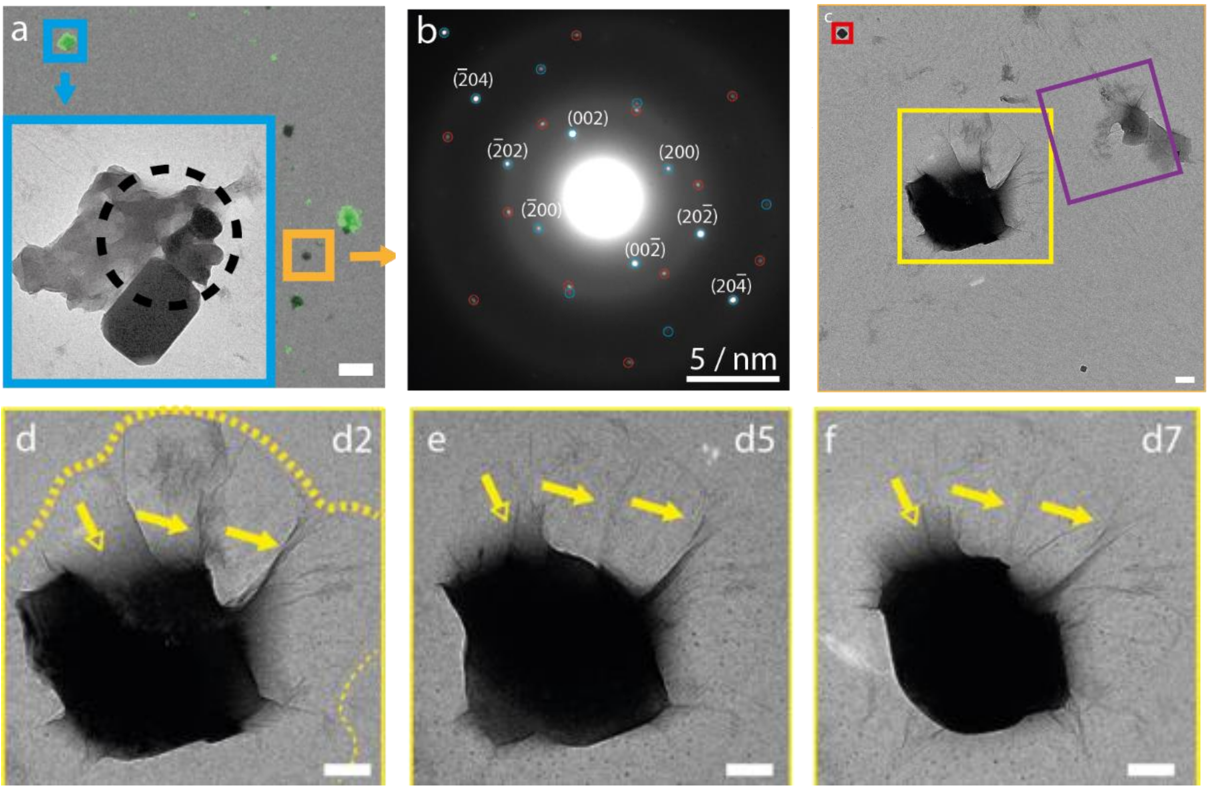

**Figure S7**. **Visualization of crystallization processes inside a GLC.** a) TEM overview imaged two days after thawing, overlaid with live-FM (green) to indicate the GLCs. Inset shows a high magnification LP-TEM image of the GLC in the blue box. b) SAED pattern taken at the position indicated by the black dashed circle in (a). Blue circles show the contribution of the NaCl crystal, and red circles the contribution of the graphene monolayer. c) Enlargement of the area marked by the orange box in (a) were multiple GLCs are present (purple and yellow boxes) and a crystal that is not encapsulated by graphene (red box). d-f) Large crystal inside a GLC imaged at multiple time points after thawing (Figure S7d: 2 days; Figure S7e: 5 days; Figure S7f: 7 days) shows morphological changes. Graphene wrinkles (close yellow arrow), outline of the GLC (dotted yellow line) and the intensity gradient of the liquid surrounding the crystal (open yellow arrow) are visible at all time points. Accumulative dose: a) 0.75e$^-$/Å$^2$; c, d) 0.15 e$^-$/Å$^2$; e) 0.30 e$^-$/Å$^2$; f) 0.45 e$^-$/Å$^2$

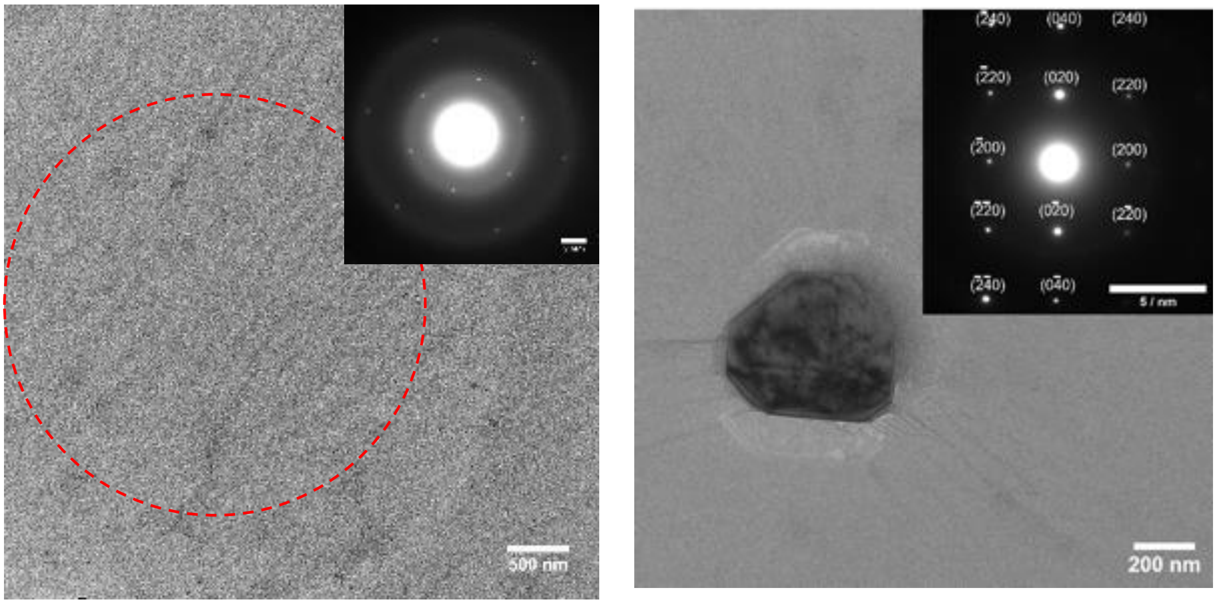


**Figure S8**: Selected Area Electron Diffraction (SAED) taken on the grid close to a liquid pocket showing the reflections corresponding to the graphene monolayer and the diffuse rings of the amorphous carbon (a) and from an encapsulated single NaCl crystal (zone axis [10-29](b).

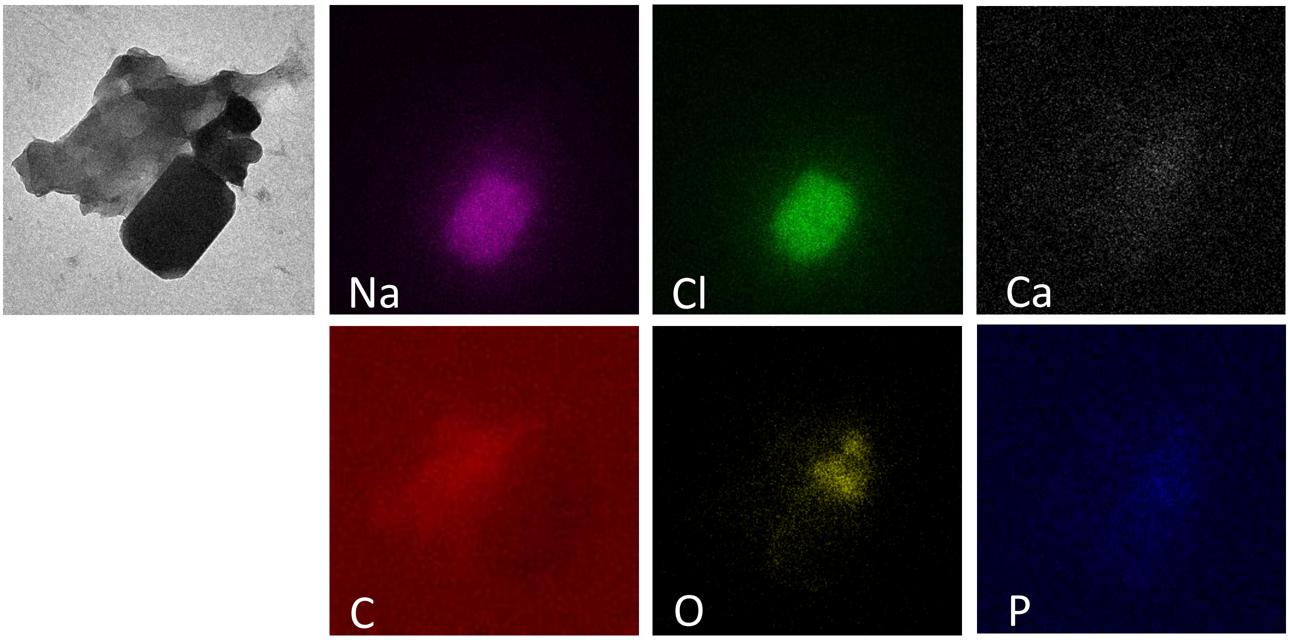


**Figure S9:** EDX map showing the composition of the shaped crystal.

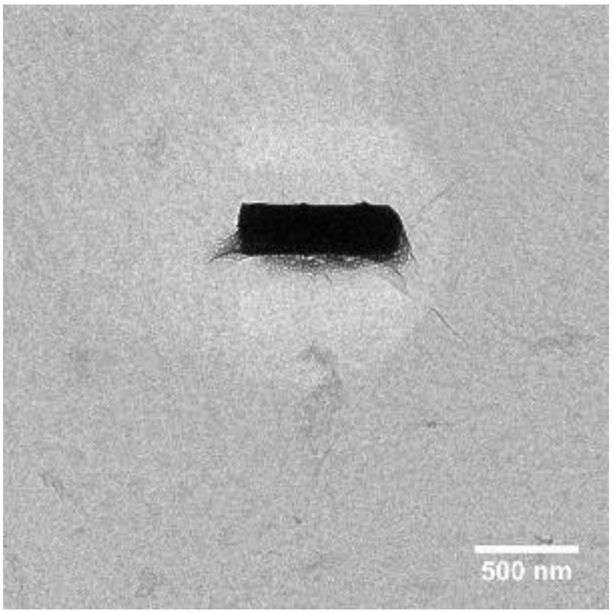


**Figure S10:** Crystal imaged using high electron dosage ($e^-$-dose > 100 e/Å$^2$). Bubbling becomes visible as a result of radiolysis.

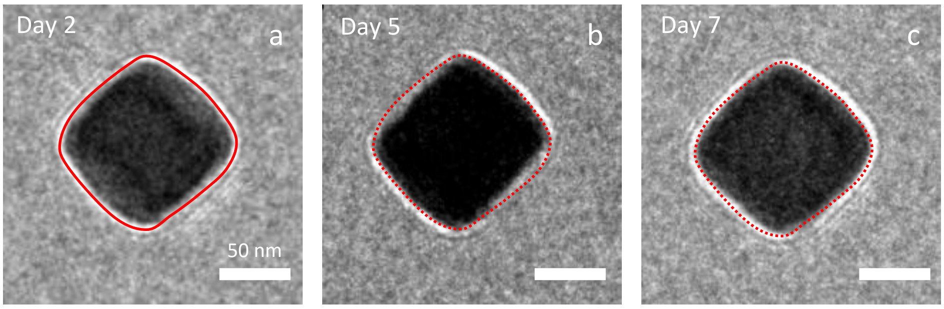


**Figure S11:** Not encapsulated crystal showing Fresnel fringes all around them. Fresnel fringes only appear at the sample edge in out of focus imaging and not when the crystal is encapsulated because the liquid will create an intensity gradient instead of a sharp edge. Besides the absence of a fluorescent signal, there are also no wrinkles, pocket outline or intensity gradient which indicates that the crystal is not encapsulated. Since there is no mineral precursors supply, these crystals do not show crystal growth, but instead retain their original external shape and size.

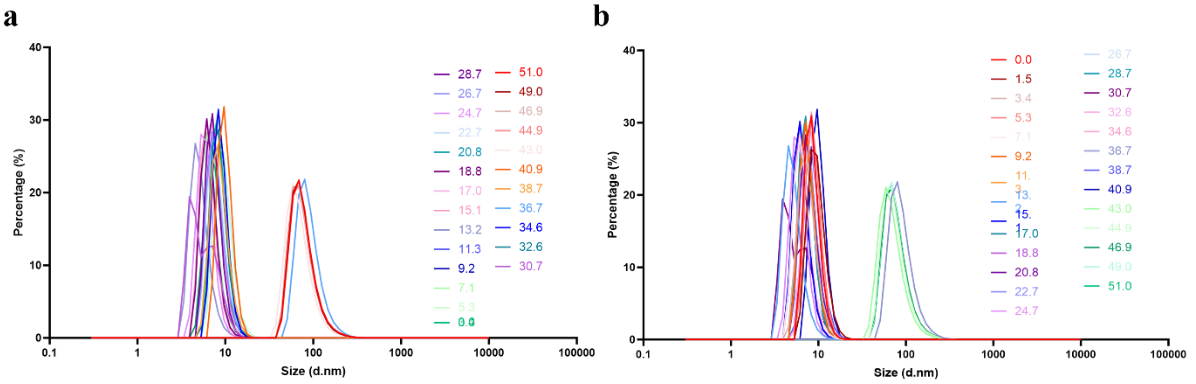


**Figure S12. Dynamic light scattering experiments of CPP formation showing the a) number and the b) intensity plots.**

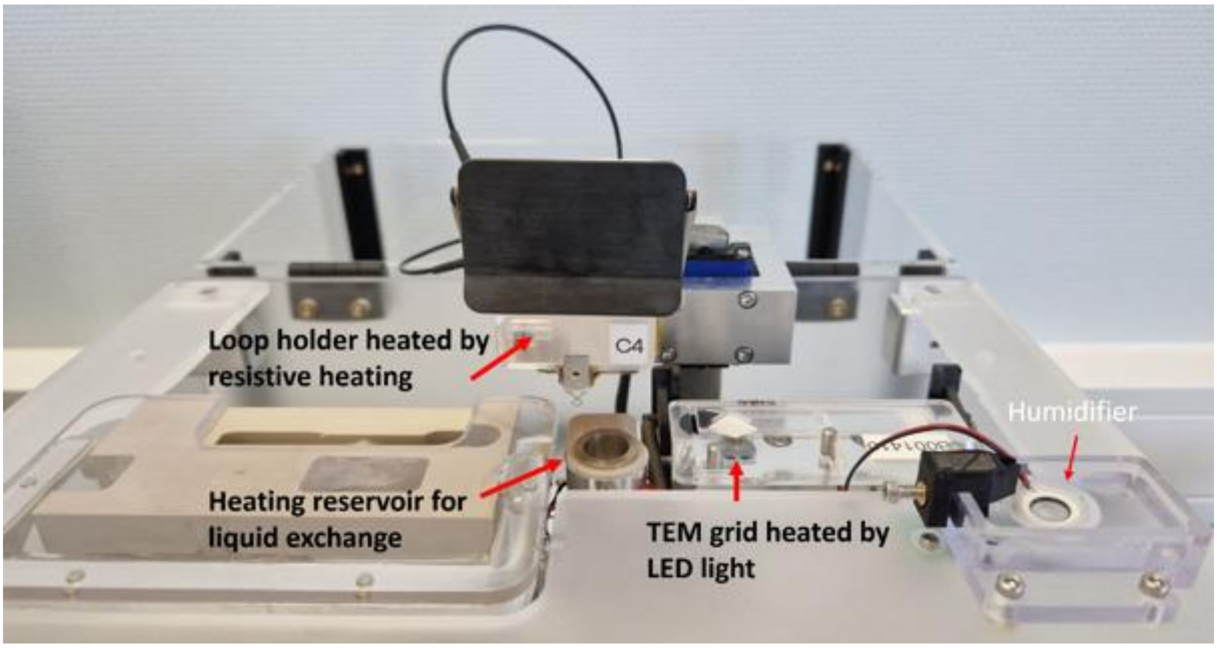


**Figure S13: NAIAD-1 Mod-T modified to have control over temperature and humidity.** The liquid in the loop is heated to 37 °C. A reservoir was added with heated (37 °C) liquid to exchange fluids in the loop. TEM grid was heated (37 °C) and humidifier to establish 96% humidity.

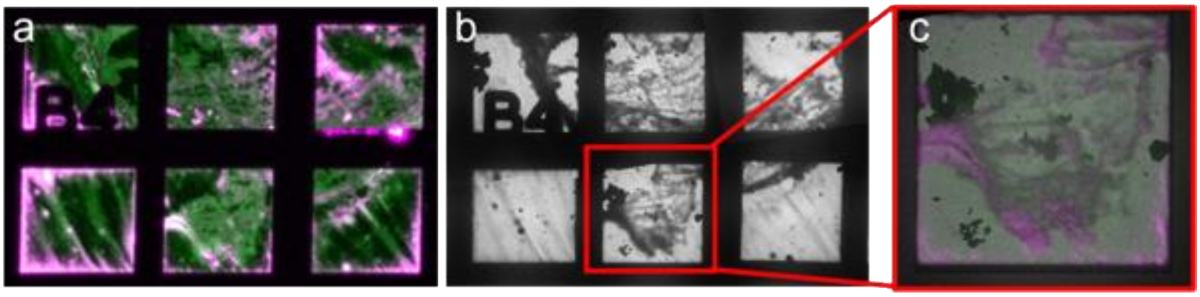


**Figure S14: Cryo-CLEM workflow to determine the regions of interest for the visualization of CPP complexation.** a) Finder grid imaged with fluorescent microscopy (CF680 for liquid in magenta and Fetuin-A in green). Overlapping signal appears in white. b) Cryo-TEM overview of the same area. c) correlation of the fluorescent and cryo-TEM image.

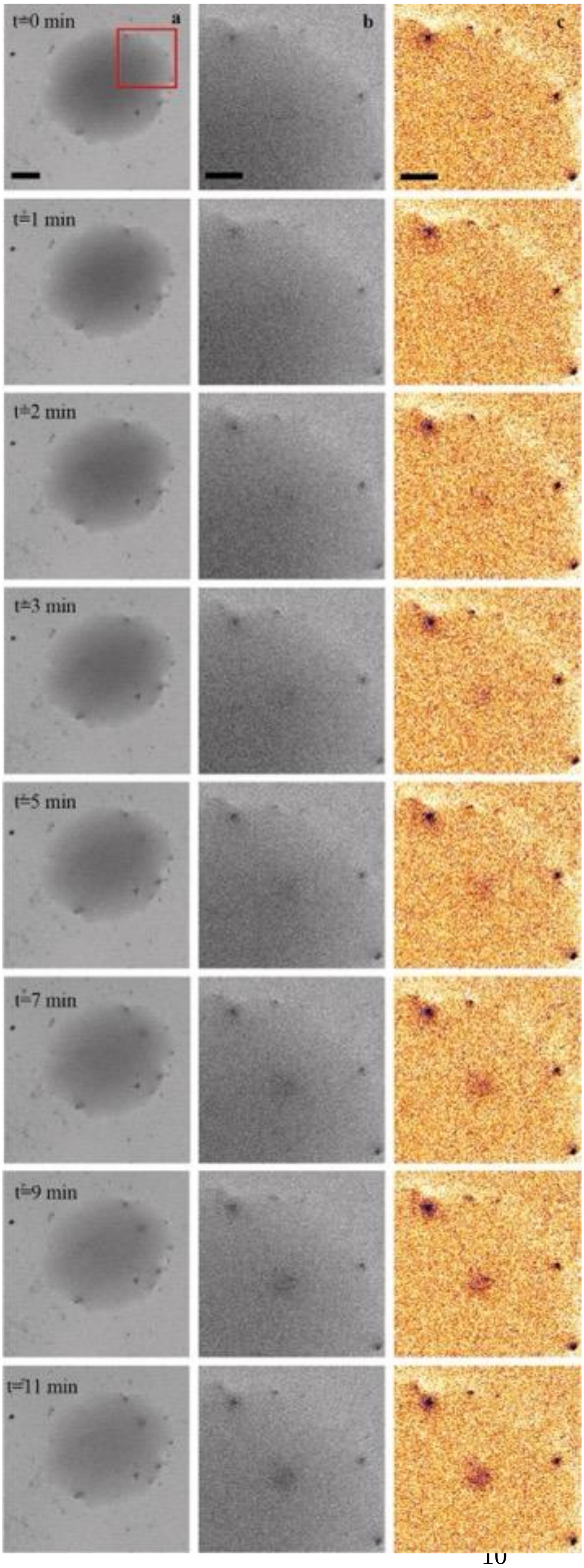

t=0 min
a
b
c
t=1 min
t=2 min
t=3 min
t=5 min
t=7 min
t=9 min
t=11 min

**Figure S15. Formation of CPP in a GLC.** a) Entire graphene liquid cell over the imaging time of 11 minutes. b) Selected region of interest (red square (a)) over the imaging time of 11 minutes. c) Same area as (b) but after contrast enhancement, denoising and shading correction in white/red contrast.
